# From electrophysiology to drink: Adolescent alcohol consumption predicted by differences in functional connectivity and neuroanatomy

**DOI:** 10.1101/2024.04.29.591615

**Authors:** Alberto del Cerro-León, Luis Fernando Antón-Toro, Danylyna Shpakivska-Bilan, Marcos Uceta, Alejandro Santos-Mayo, Pablo Cuesta, Ricardo Bruña, Luis M García-Moreno, Fernando Maestú

**Affiliations:** Center of Cognitive and Computational Neuroscience, Universidad Complutense de Madrid (UCM), Madrid, Spain; Department of Experimental Psychology, Cognitive Processes and Speech Therapy, Universidad Complutense de Madrid (UCM), Madrid, Spain; Department of Psychology, University Camilo José Cela (UCJC), Madrid, Spain; Department of Cellular Biology, Faculty of Biology, Complutense University of Madrid (UCM), Madrid, Spain; Health Research Institute of the Hospital Clínico San Carlos (IdISSC), Madrid 28040, Spain; Department of Radiology, Complutense University of Madrid, Madrid, Spain; Department of Psychobiology and Methodology in Behavioral Science, Faculty of Education, Complutense University of Madrid (UCM), Madrid, Spain

**Keywords:** Electrophysiology, Alcohol, Functional connectivity, Magnetic resonance imaging, MEG

## Abstract

Alcohol consumption during adolescence has been associated with neuroanatomical abnormalities and the appearance of future disorders. However, the latest advances in this field point to the existence of risk profiles which may lead to some individuals into an early consumption. To date, some studies have established predictive models of consumption based on sociodemographic, behavioural, and anatomical-functional variables using MRI. However, the neuroimaging variables employed are usually restricted to local and hemodynamic phenomena. Given the potential of connectome approaches, and the high temporal dynamics of electrophysiology, we decided to explore the relationship between future alcohol consumption and electrophysiological connectivity measured by MEG in a cohort of 83 individuals aged 14 to 16. We calculated predictive models throughout multiple linear regressions based on behavioural, anatomical, and functional connectivity variables. As a result, we found a positive correlation between alcohol consumption and the functional connectivity in frontal, parietal, and frontoparietal connections. Also, we identified negative relationships of alcohol consumption with neuroanatomical variables. Finally, the linear regression analysis determined the importance of anatomical and functional variables in the prediction of alcohol consumption but failed to find associations with impulsivity, sensation-seeking, and executive function scales. As conclusion, the predictive traits obtained in these models were closely associated with changes occurring during neurodevelopment, suggesting the existence of different paths in neurodevelopment that have the potential to influence adolescents’ relationship with alcohol consumption.

**Significance statement:** To understand the onset of heavy drinking habits and develop prevention strategies, we need to characterize predisposition profiles at early ages. This longitudinal work provides important evidence by showing how adolescents at risk for engaging in alcohol behaviors showed resting-state functional connectivity and grey matter differences years before. The combination of these metrics allows us to establish predictive models of future alcohol episodes. In addition, differences in functional connectivity showed a positive relationship with behavioral variables such as lower executive functions and higher on the sensation-seeking. These predisposition phenotypes may rely on divergent neurodevelopmental pathways and deeper neurobiological abnormalities, such as dysfunctions of inhibitory neurotransmission and/or a genetic background of vulnerability.

## 1 Introduction

Alcohol consumption is highly tolerated in our society, especially among young people and adolescents, a sector of the population that is increasingly normalizing its consumption as a form of recreation ^1^. Such consumption typically begins at the age of 14 and intensifies throughout adolescence until it reaches episodes of intensive consumption at the age of 16 ^2^. Unfortunately, since adolescence is a maturational stage, during this period the brain becomes particularly vulnerable to external agents ^3^. In this regard, numerous studies have described how alcohol alters brain neurodevelopment at molecular ^4^, structural ^5,6^ and functional level ^7,8^. Consequently, at the behavioural level, alcohol drinking during adolescence has been associated with deficits in executive function ^9^, impulsivity ^10^, and emotional regulation ^11^. Nevertheless, it’s important to recognize that cross-sectional studies on alcohol consumption can’t definitively determine if impairments are caused by alcohol use or by pre-existing brain conditions that lead to risky behaviours ^12^. Recent longitudinal studies with a prospective experimental design have identified behavioural, anatomical and functional differences in young individuals who later engage in alcohol use. On the one hand, it has been observed that adolescents prone to alcohol habits reflect high scores in sensation seeking and deficits in executive functions, with a special involvement of inhibitory control and executive memory ^5^. Likewise, structural MRI studies have described how deviations in subcortical volumes such as cerebellum and caudate nucleus ^13^ and smaller grey matter and greater white matter volumes in frontal and temporal regions ^14^ are associated with higher levels of future consumption. Finally, fMRI studies enlightened that adolescents prone to alcohol abuse show blunted BOLD activation during inhibition, working memory ^5^ and rest ^15^.

Despite the relevance of these results, the study of the integrity of brain’s functional networks has become crucial to the understanding of the neural mechanisms associated with complex behaviours such as substances use. Functional connectivity (FC) is defined as the statistical dependence between the activity of two or more brain regions and allows us to quantify the connections based on the temporal relationship in their physiological signals ^16^. To study the phenomenon of functional networks, traditional approaches have used resting-state paradigms that measure the synchronization of spontaneous oscillatory activity when participants are not engaged in any specific task ^17,18^. Using these metrics, some fMRI studies found that lower FC between prefrontal cortices and subcortical regions implicated in the motivated and reward-directed behaviour are predictors of alcohol consumption ^19^. However, in spite of the excellent spatial resolution of fMRI, many of the functional processes of the brain occur at very short time scales that only electrophysiology techniques (M/EEG) can cover ^18,20^. To our knowledge, only one study have examined the relationship between M/EEG FC and alcohol rates during adolescence, depicting a link between alpha, beta and gamma hyperconnectivity and the development of binge drinking profiles also associated with sensation seeking and impaired executive functioning ^21^. Moreover, none of the current studies have integrated M/EEG functional connectivity metrics in predictive analysis with other variables from different modalities. The present study aims to depict predisposition profiles towards alcohol consumption based on functional brain connectivity, structural (grey and white matter volumes), and behavioural (executive function, impulsivity and sensation seeking) variables throughout multiple linear regressions analysis. According with previous results, we expect that higher FC, reduced cortical volumes, and worse behavioural profiles would be predictive of higher alcohol use during later stages.

## 2 Materials & Methods

### 2.1 Participants

The sample of this study was constituted by adolescents recruited from different high schools within the community of Madrid within the framework of two longitudinal projects financed by the Spanish Ministry of Health. Both projects followed the same evaluation protocol, which consisted in two stages separated by a follow-up period of two years. Prior to the experiment, all participants indicated no history of alcohol consumption, family alcohol use disorders and absence of psychiatric or neurological disorders. Additionally, all subjects fulfilled the Alcohol Use Disorder Identification Test (AUDIT) ^22^ and completed a semi-structured interview about their drug use habits in order to discard any potential consumer at this stage. In the first stage (before alcohol use onset), 148 adolescents accepted to participate in the Magnetoencephalography (MEG) study which consisted of a 5 minutes eyed-closed resting-state, and 121 of them also participated in the MRI study. Additionally, all participants fulfilled neuropsychological and behavioural self-reported questionaries to measure the dysexecutive behaviour, impulsivity and sensation seeking variables. After a follow-up period of two years, 92 participants underwent again the AUDIT test and a semi-structured interview. Using this information, we calculated the quantity of Standard Alcohol Units (SAUs) consumed during regular drinking episodes for each participant. This calculation considered the number of beverages consumed within a 2–3-hour period, as well as the specific type of beverage. Tobacco and cannabis use were controlled discarding those participants with a regular use of these substances. Finally, after quality control of the MEG and MRI data, a final sample of 83 subjects (mean age 14.42 ± 0.60, 41 males and 42 females) completed the whole protocol and was selected for the study. Informed consent was obtained from all participants and their parents or legal guardians following the guidelines outlined in the declaration of Helsinki. The ethical committee of the Universidad Complutense de Madrid granted approval for the study.

### 2.2 MRI recordings and volumetry

The participants underwent a 3D T1-weighted high-resolution brain MRI scan with a power of 1.5 T from the Santa Elena Foundation (General Electric Optima MR450w, echo time = 4.2 ms, repetition time = 11.2 ms, inversion time = 450 ms, flip angle = 12°, field of view = 100, acquisition matrix = 256 × 256, and slice thickness = 1 mm) or the Clinical Hospital of Madrid (General Electric Signa HDxt, echo time = 4.2 ms, repetition time = 11.2 ms, inversion time = 450 ms, flip angle = 20°, field of view = 100, acquisition matrix = 256 × 256, and slice thickness = 1 mm).

MRI images from each participant were processed with Freesurfer image analysis suite (version 5.3.0), which is documented and freely available for download online (http://surfer.nmr.mgh.harvard.edu/). To extract subcortical volumes (Nucleus accumbens (NAcc), caudate, putamen and grey and white matter from cerebellum) we used the automatic segmentation based on probabilistic information estimated from a set of 40 training segmentations, manually labelled using the Center for Morphometric Analysis conventions ^23^. Additionally, a surface atlas of brain lobes made with the tksurfer tool was used to calculate the values of grey matter and white matter of the cortex following a lobe parcellation. All the volumes obtained from these methods were eventually corrected for the total brain volume of each participant.

### 2.3 Neuropsychological scales

Widely used self-report scales were used to assess traits of impulsivity, sensation seeking and dysexecutive behaviours. Barrat impulsivity scale (BIS-11) ^24^ was used to measure impulsivity traits. Sensation-seeking scale (SSS-V) ^25^ was used to assess thrill and adventure seeking, disinhibition, experience seeking and susceptibility to boredom, that reflect the tendency to experience risky behaviours. Three complementary scales were used to assess executive dysfunction: Barkley deficits of executive function scale in its abbreviated form (BDEFS) ^26^, behavioural rating inventory of executive function (BRIEF) ^27^ and dysexecutive questionnaire (DEX) ^28^. BRIEF test was corrected for the negativity and sincerity subscales by detecting outliers as those subjects who scored above 3 standard deviations from the mean. Due to the lack of normality in this test, the outlier scores were replaced by the median value. In order to reduce the dimensionality of the executive questionnaires, a principal component analysis was performed using the BRIEF global executive index and the BDEFS and DEX scores.

### 2.4 MEG recordings and signal processing

MEG data were collected using a 306-channel Elekta Neuromag system, placed in a magnetically shielded room at the Center for Biomedical Technology in Madrid, Spain. During data acquisition, an online fir type anti-alias filter was applied with a frequency range of 0.1 to 330 Hz, and a sampling rate of 1,000 Hz was used. To minimize environmental noise and correct subject movements, a temporal extension of the signal space separation method (correlation window of 10 s, and correlation limit of 0.9) was employed ^29^. Afterwards, Fieldtrip software ^30^ was used on Matlab R2020b to automatically detect artifacts in the signal that were visually confirmed by an MEG expert. Eventually, artifact-free data were segmented in epochs of 4 seconds plus 2 seconds of real data at each side as padding.

### 2.5 Source-space reconstruction

Individual MEG signals were estimated at the source level using the subject’s own T1-weighted MRI. A homogeneous grid of source positions, labeled according to the automated anatomical labeling (AAL) atlas ^31^, was employed as the source model. These 1202 positions, labeled as 78 cortical regions of interest (ROIs), were then transformed to the subject’s space using a linear transformation between the MNI template and the participant’s T1-weighted MRI. Additionally, the T1 image was used to create a single-shell head model based on the inner surface of the skull ^32^. By combining the head model, source model, and sensor definition, a leadfield was constructed using a modified spherical solution. Finally, as an inverse method, we used a linearly constrained minimum variance (LCMV) beamformer.

### 2.6 Functional connectivity analysis (FC)

Functional connectivity (FC) was assessed using the phase locking value (PLV), a measure of phase synchronization that examines the phase differences between two time series. To calculate the PLV for each frequency band and epoch, we use the procedure followed by Bruña et al., 2018. Consequently, this algorithm yielded symmetrical whole-brain matrices with dimensions of 1202 × 1202 nodes for each participant and frequency band (theta: 4-8 Hz, alpha: 8-12 Hz, low-beta: 12-20 Hz, high-beta: 20-30 Hz and gamma: 30-45 Hz). For each pair of the 78 cortical regions, the PLV value was calculated using the root-mean-square of PLV values connecting these ROIs in a 78 by 78 whole brain FC matrix. For a detailed description of the MEG recordings, processing, source reconstruction and PLV calculations see Supplementary Materials.

### 2.7 Statistical analysis

#### 2.7.1 Cluster based permutation test (CBPT)

The relationship between the PLV values of each ROI pair and the SAUs was assessed by a CBPT analysis separately for each frequency band ^33^. Due to the results obtained previously ^21^, in which positive relationships were found between future consumption levels and functional connectivity values, 1-tailed spearman partial correlations were performed under the assumption that the results should have a positive relationship. Partial correlations were controlled by age, sex, and data sample. We defined a cluster as a set of spatially adjacent links that presented a significant partial correlation (only links with a correlation significance less than 0.005 were considered) in the same direction between the PLV values and SAUs. Only clusters including at least 1% of the grid (i.e., a minimum of 12 nodes) were considered. The Spearman rho values were transformed into Fisher Z values, and the cluster-mass statistics were computed as the sum of the Z values of all links within the cluster. The p value for each cluster was calculated in a nonparametric fashion, using a null distribution generated by the mass of the main cluster obtained over 50000 random permutations (shuffled versions) of the data ^34^. Only those clusters that resulted significant (p< 0.05) after this step were considered in further analyses. Then, we used the average of the PLV values of the members to obtain a representative PLV of the cluster. Of note, this FC marker would be indicating that the PLV of the clusters appeared to be associated with future alcohol consumption levels. Finally, to observe the relationship of these networks with the behavioral scales, similar CBPT analysis was elaborated considering only the connectivity matrices resulting from the previous analysis and the scores of SSS-V, BIS-11 and the first component of the executive questionnaires (only links with a correlation significance less than 0.01 were considered). Since electrophysiological signals obtained with EEG or MEG are commonly affected by volume conduction and leakage effects, all links obtained in the CBPT analysis were subsequently corrected using the correlation between spatial filters of the beamformer and power in the band of interest of the ROIs involved as covariates. Only the links that maintained significance under the selected significance level in each analysis were considered to surpass this correction.

#### 2.7.2 Multiple regression analysis

In the case of the behavioral analysis, scores obtained in the self-reported scales (BIS-11 and SSS-V) and first component of the executive questionnaires were considered as potential predictors in a stepwise multiple regression model. Neuroanatomical variables were grouped into predictive models of gray matter (frontal, temporal and parietal lobes, cerebellum, nucleus accumbens, caudate and putamen) and white matter (frontal, temporal and parietal lobes and cerebellum) volume. Finally, the predictors obtained in the previous models were employed as independent variables along with the FC results of the exploratory analysis in a multimodal regression analysis. All these analyses were performed on the SPSS software using the consumption ratio (number of SAUs) as the dependent variable.

The leave-one-out method (LOOCV) was used as the cross-validation technique, in which regression models in behaviour, volumetrics, and multimodal data were performed as many times as there were subjects in the sample. In each iteration, a different subject was excluded and used as the test set to predict their consumption value. Finally, the actual and predicted SAU values from the different models were used to calculate the root mean squared error and the R-squared adjustment. Additionally, those predictors that were selected in all multimodal models were used in the entire sample to extract the corresponding coefficients.

## 3 Results

During the follow-up phase, the participants exhibited distinct consumption patterns from abstinence to heavy drinking, with a mean consumption of 3.74 ± 2.81 SAUs. To elucidate the origins of these disparities, comprehensive analyses from varied methodologies were conducted, facilitating the identification of preliminary biomarkers associated with alcohol consumption risk during adolescence. Predictive models were formulated using behavioural and volumetric data, selecting independent variables previously established to be correlated with adolescent alcohol consumption in extant literature. Nonetheless, concerning electrophysiology, a preliminary exploration of the association between alcohol consumption and functional connectivity was undertaken prior to the predictive analysis, given the limited literature available on this topic.

### 3.1 Characterization of the relationship between FC and SAUs

To describe the relationship between functional connectivity and future alcohol consumption, an exploratory analysis in a full-head model based on AAL atlas was performed. The CBPT analysis revealed 5 significant networks in the theta, alpha, low beta, high beta and gamma bands, with positive correlations between PLV and future SAUs. The number of links that surpass the volume conduction and leakage correction are indicated in parentheses next to the value of links in the initial analysis. Theta network (Figure 1) consisted of 64 (57) links between bilateral prefrontal, frontotemporal, parietal, and left occipital regions with a higher implication of the left frontal lobe (*p* < 0.05, rho = 0.49). In the alpha band (Figure 2), the network presented 114 (107) links distributed in intra- and interhemispheric connections between frontal, parietal and temporal pole regions (*p* = 0.03, rho = 0.42). Results in the low beta band (Figure 3) presented a network with 65 (52) links spanning frontotemporal and occipital intra- and interhemispheric connections (*p* < 0.05, rho = 0.46). Meanwhile, the high beta band results (Figure 4) showed a network with 122 (95) links with a wide distribution between fronto-temporal and parieto-occipital regions, observing a high predominance of connections with the left hemisphere (*p* = 0.02, rho = 0.52). Finally, the gamma band results (Figure 5) described a network consisting of 104 (77) connections involving frontotemporal and parieto-occipital regions with a high predominance of connections to the left hemisphere (*p* = 0.03, rho = 0.47). Figures 1 to 5 show the cortical distribution of the significant links, as well as the correlation between the mean PLV values of the network and the SAUs values. Additionally, the networks obtained in the exploratory analysis were evaluated in each of the datasets separately. Apart from the results in the low beta band, this analysis showed that the networks obtained have significant positive correlations with future consumption and similar behaviour regardless of the sample.

**Figure 1.**
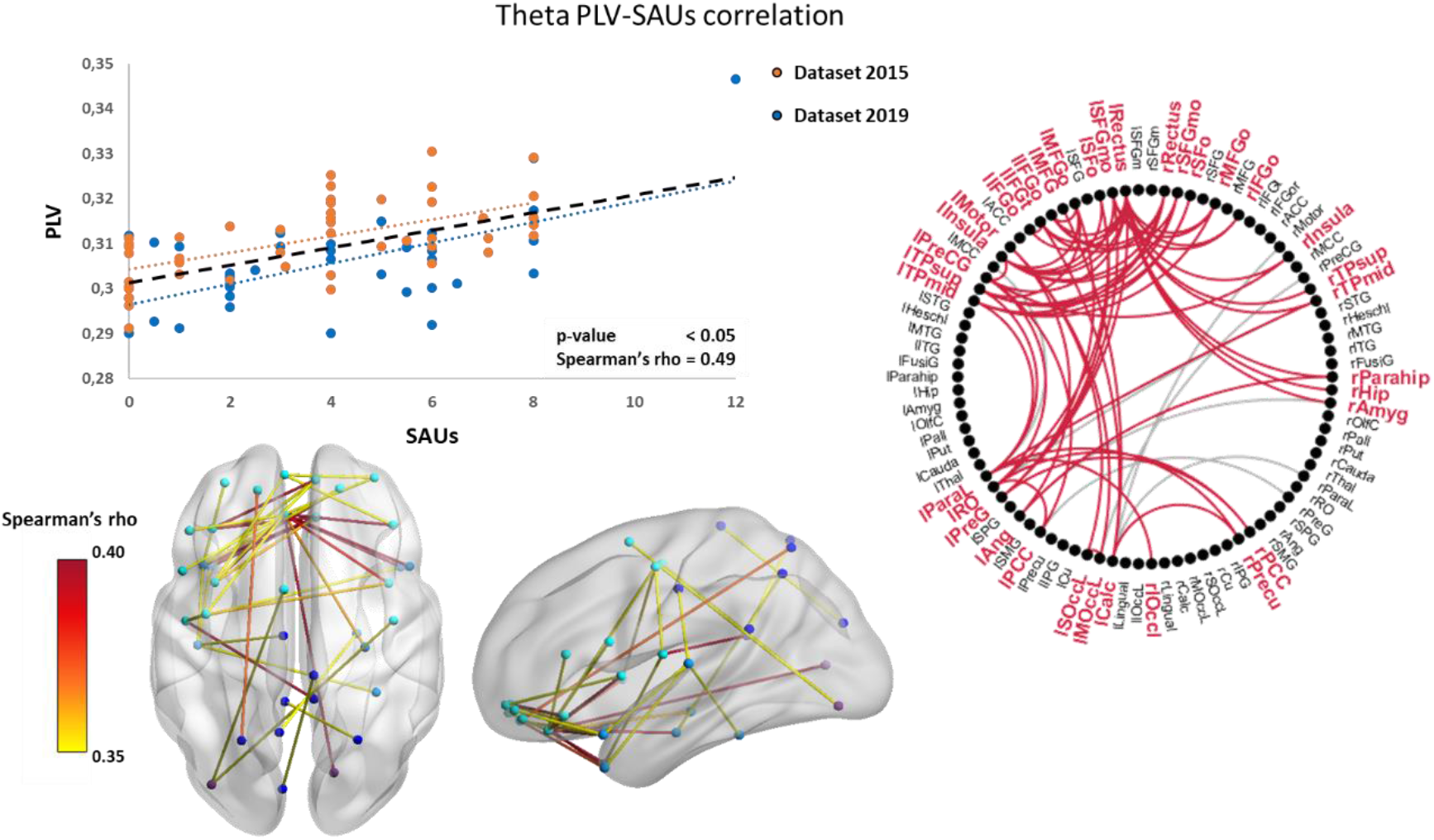
Correlation of functional connectivity in the Theta band (PLV) with future consumption levels (SAUs). The orange and blue dots represent the individual values from the 2015 and 2019 databases respectively. The black dashed line represents the trend line for the adolescent population as a whole. At the bottom, the distribution of links with a rho greater than 0.35 on a template brain in horizontal and sagittal view is depicted. The yellow-red gradation represents the rho value of each individual link. On the right there is a summary graph of the spatial distribution of the network. The grey lines represent those links that did not surpass the volume conduction and leakage correction.

**Figure 2.**
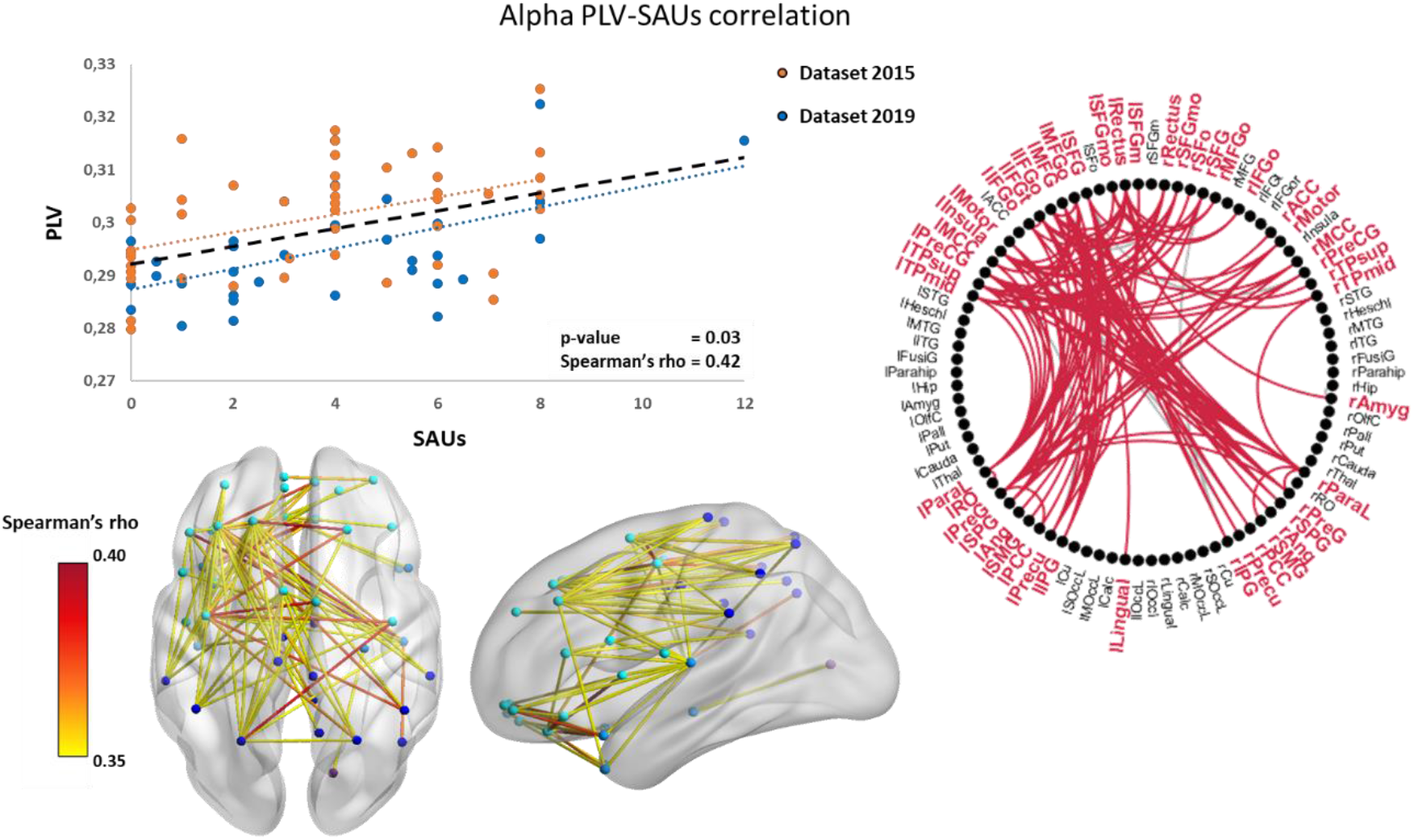
Correlation of functional connectivity in the Alpha band (PLV) with future consumption levels (SAUs). The orange and blue dots represent the individual values from the 2015 and 2019 databases respectively. The black dashed line represents the trend line for the adolescent population as a whole. At the bottom, the distribution of links with a rho greater than 0.35 on a template brain in horizontal and sagittal view is depicted. The yellow-red gradation represents the rho value of each individual link. On the right there is a summary graph of the spatial distribution of the network. The grey lines represent those links that did not surpass the volume conduction and leakage correction.

**Figure 3.**
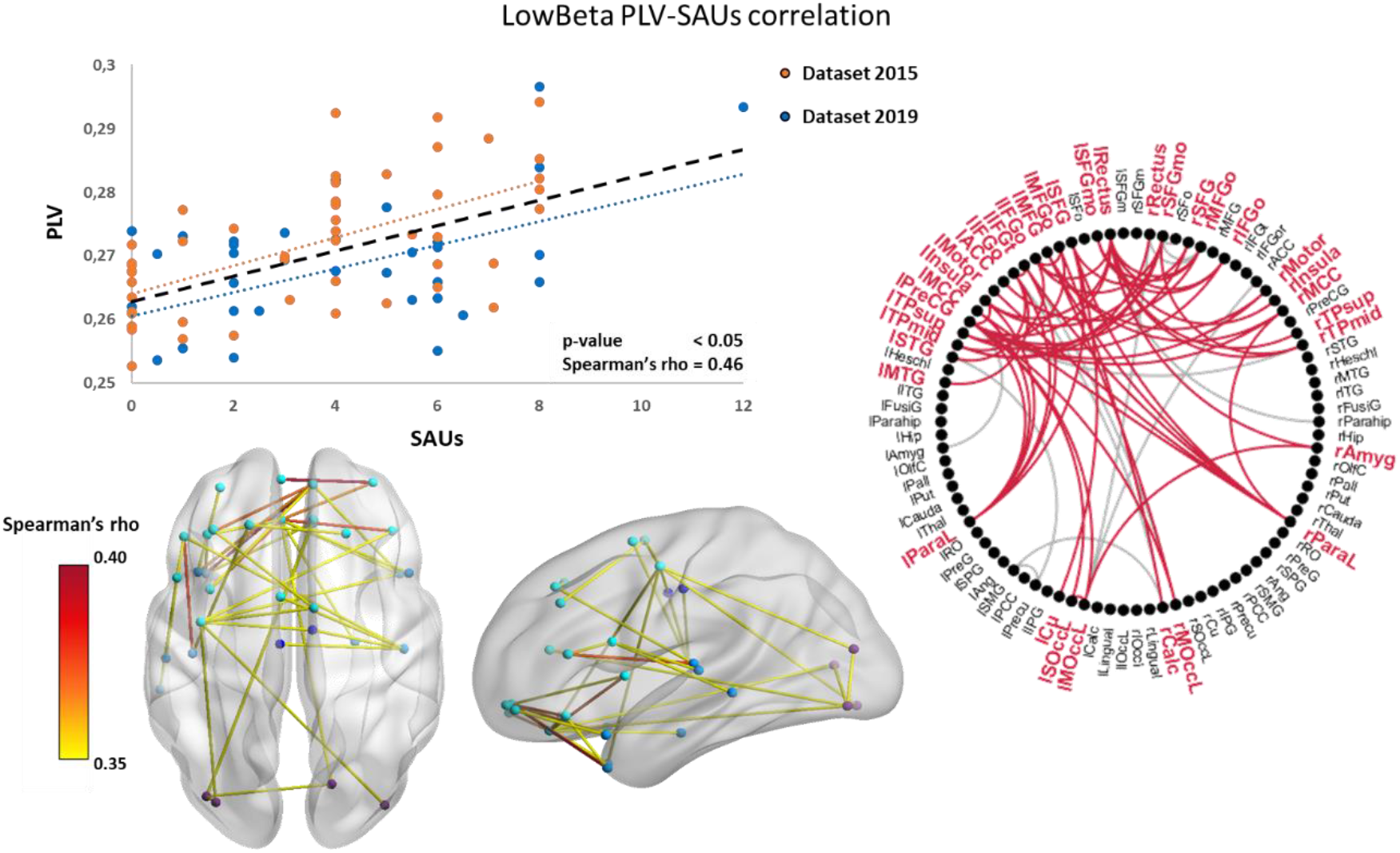
Correlation of functional connectivity in the Low Beta band (PLV) with future consumption levels (SAUs). The orange and blue dots represent the individual values from the 2015 and 2019 databases respectively. The black dashed line represents the trend line for the adolescent population. At the bottom, the distribution of links with a rho greater than 0.35 on a template brain in horizontal and sagittal view is depicted. The yellow-red gradation represents the rho value of each individual link. On the right there is a summary graph of the spatial distribution of the network. The grey lines represent those links that did not surpass the volume conduction and leakage correction.

**Figure 4.**
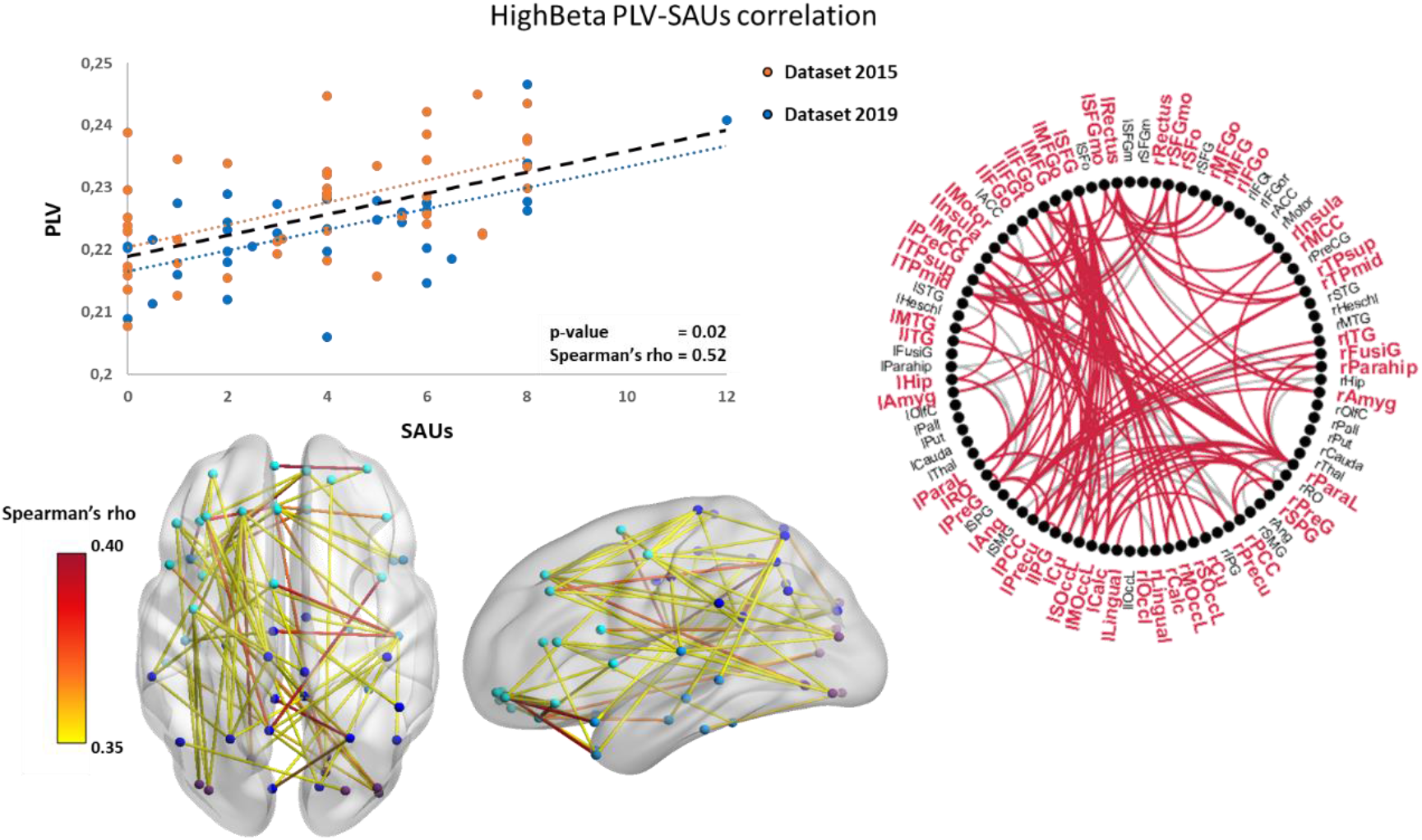
Correlation of functional connectivity in the High Beta band (PLV) with future consumption levels (SAUs). The orange and blue dots represent the individual values from the 2015 and 2019 databases respectively. The black dashed line represents the trend line for the adolescent population. At the bottom, the distribution of links with a rho greater than 0.35 on a template brain in horizontal and sagittal view is depicted. The yellow-red gradation represents the rho value of each individual link. On the right there is a summary graph of the spatial distribution of the network. The grey lines represent those links that did not surpass the volume conduction and leakage correction.

**Figure 5.**
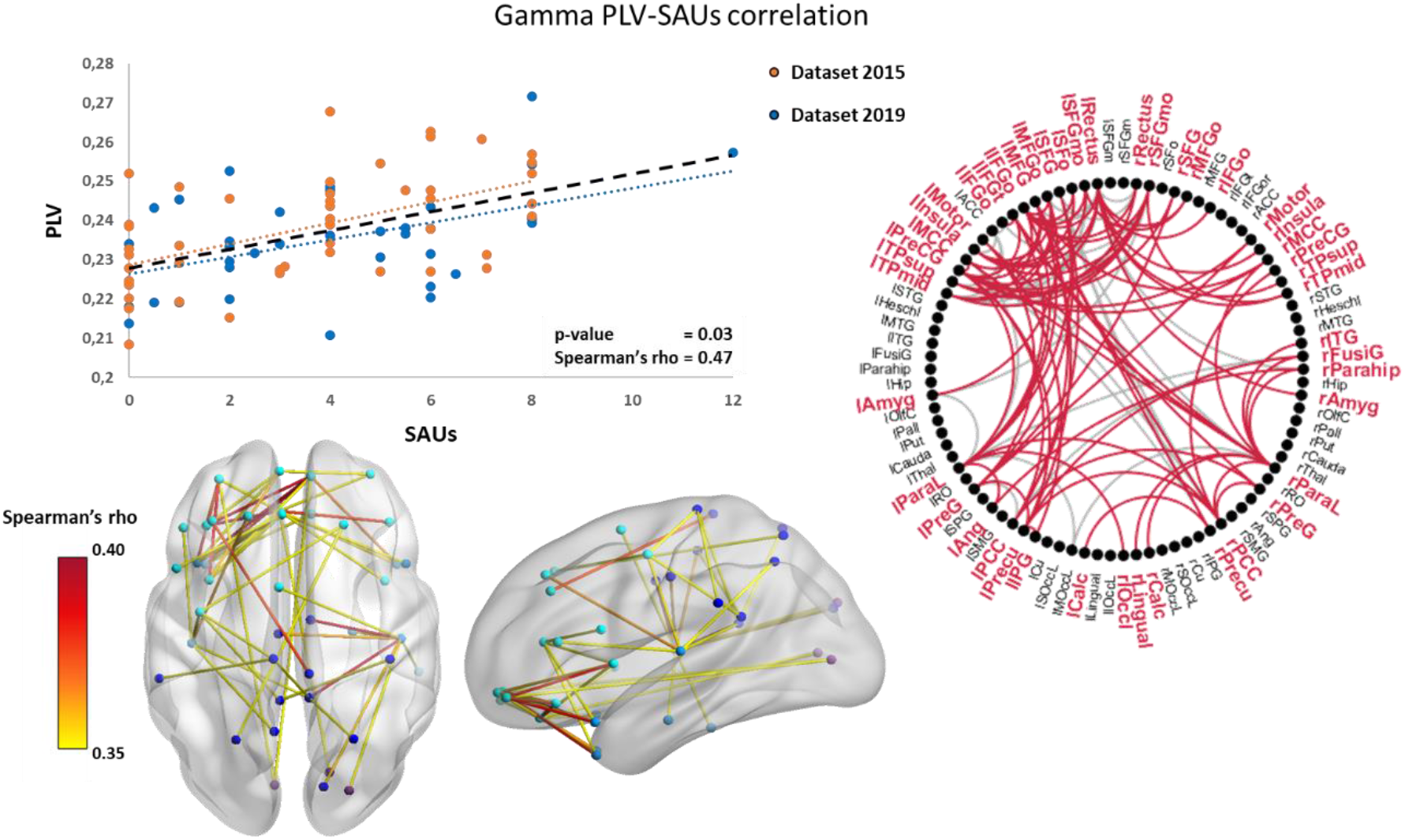
Correlation of functional connectivity in the Gamma band (PLV) with future consumption levels (SAUs). The orange and blue dots represent the individual values from the 2015 and 2019 databases respectively. The black dashed line represents the trend line for the adolescent population. At the bottom, the distribution of links with a rho greater than 0.35 on a template brain in horizontal and sagittal view is depicted. The yellow-red gradation represents the rho value of each individual link. On the right there is a summary graph of the spatial distribution of the network. The grey lines represent those links that did not surpass the volume conduction and leakage correction.

**Figure 6.**
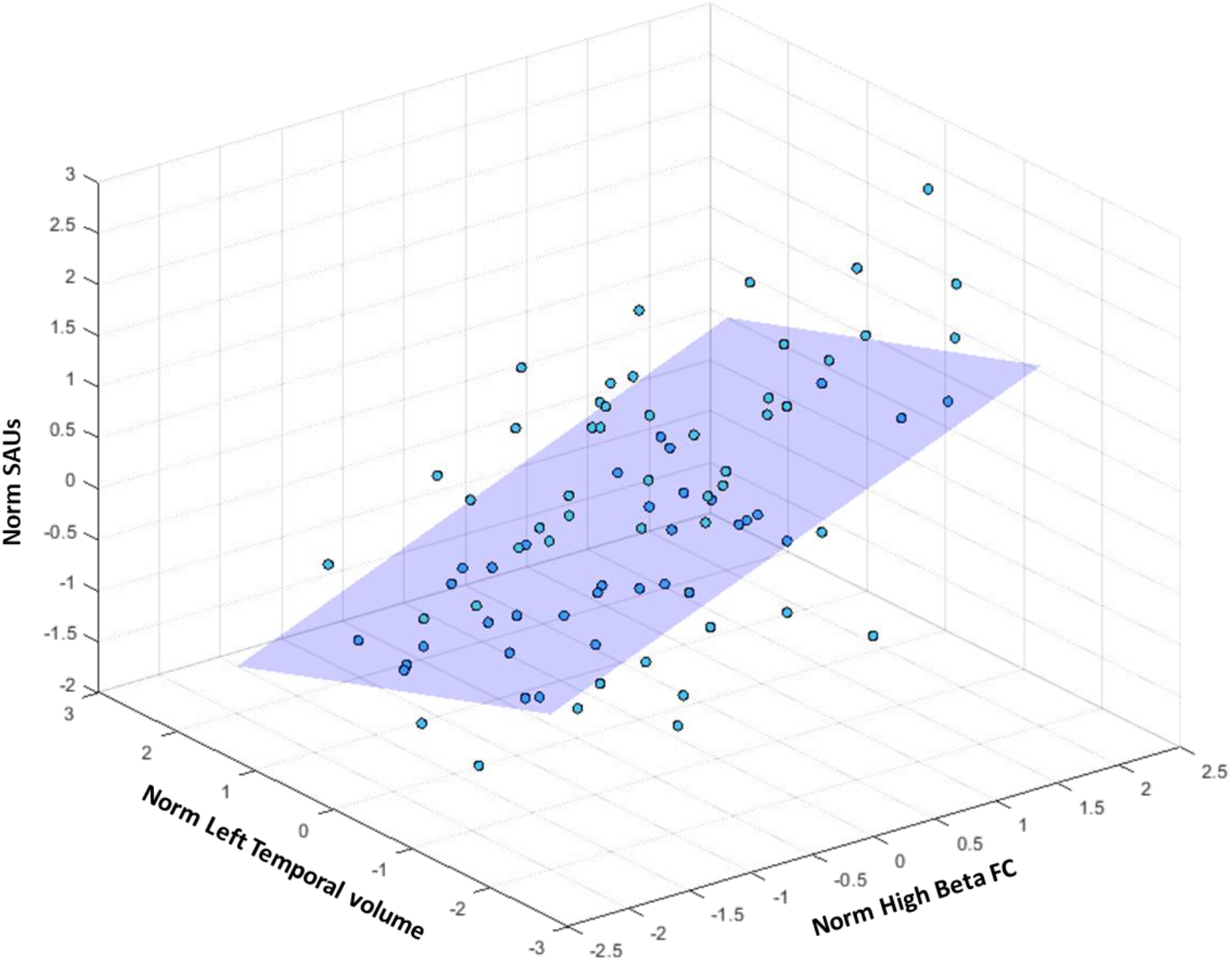
Multiple linear regression model with left temporal cortex and high beta functional connectivity as predictors. The individual points represent the values obtained in the analyzed sample. The plane represents the values predicted by the regression model. The normalized value of functional connectivity in the high beta band is plotted in the x-coordinates. The normalized volume of the left temporal cortex is plotted in the y-coordinates. The normalized values of alcohol consumption levels are plotted in z-coordinates.

### 3.2 Stepwise multiple linear regression analysis

After identifying networks that exhibited a significant correlation with prospective alcohol consumption, predictive models were developed using both grey and white matter volumetric measurements, as well as behavioural parameters. Subsequently, to merge the findings from various modalities, a multivariate linear regression analysis was conducted. In this model, SAUs were the dependent variable, and significant predictors identified in earlier analyses, alongside functional connectivity values, were considered as potential predictors. To provide robustness to the model, the leave one out cross validation method was used to predict the consumption values of each of the participants based on the predictors obtained in the rest of the sample. The predictors selected in each of the models are summarized in Table 1 indicating the proportion of models in which they were selected.

**Table 1.** Leave one out multiple linear regression analyses for variables from behavioural, structural and functional neuroimaging

| Predictors | Consistency (%) |
| --- | --- |
| **Behavioural Models** |  |
| **GM Models** |  |
| *Left temporal lobe ctx* | 100 |
| *Right temporal lobe ctx* | 57 |
| *Caudate* | 48 |
| **WM Models** |  |
| *Cerebellum wm* | 98 |
| *Right Frontal wm* | 98 |
| **Multimodal Models** |  |
| *High Beta FC* | 100 |
| *Left temporal lobe ctx* | 100 |
| *Cerebellum wm* | 27 |
| *Caudate* | 8 |
| *Right temporal lobe ctx* | 1 |
| *Right Frontal wm* | 1 |
| **SMSE** | **R<sup>2</sup>** |
| 2.61 | 0.54 |

*Note.* The values of consistency were calculated as the number of times that predictor appeared in the models in relation to the total number of models. RMSE: Root Mean Square Error

#### Behavioural models

The principal component analysis of the executive questionnaires (BRIEF, BDEFS and DEX) was able to extract an executive component that explained 83.9% of the variance of the questionnaires. In the regression model, we used SAUs as dependent variable, and the executive component, the impulsivity and sensation seeking scores as predictors. Results did not offer significant relationships between the selected questionnaires and future alcohol consumption levels in any of the models tested.

#### Grey matter models

The volume of the grey matter in the left frontal, right frontal, right temporal, left temporal, cerebellum, putamen, caudate and nucleus accumbens were selected as potential predictors of future SAUs. As a result of these analyses, the left temporal cortex was selected in all LOOCV models, followed by the right temporal cortex (57%) and the caudate (48%). Regarding the direction of results, the volume of the left temporal cortex and the caudate maintained a negative relationship with future consumption in comparison with the right temporal cortex was positive related with future SAUs.

#### White matter models

The multiple linear regression analysis of white matter volumetry variables (left frontal, right frontal, right temporal, left temporal and cerebellum) revealed that the volumes from the cerebellum and the right frontal lobe were selected as predictor in the 98% of the models with a negative and positive relationship respectively with future SAUs.

#### Multimodal models

Based on the predictors obtained from the structural models (GM and WM volumes) and the functional connectivity, we conducted an integrative model with the different variables. The resulting models selected high beta band functional connectivity and left temporal cortex volume as predictors of future consumption in all models tested and occasionally together with white matter volume in the cerebellum (27%), caudate (8%), left temporal cortex (1%) and right frontal white matter (1%). In parallel, the LOOCV predictions obtained a root mean squared error of ± 2.61 SAUs and an r2 fit of 0.54. Finally, the significant predictors in 100% of the models tested were evaluated in the whole sample to extract the corresponding coefficients. The results obtained in this regression are detailed in Table 2.

**Table 2.** Multiple linear regression analysis for variables from structural and functional neuroimaging

| Predictor | $\beta$ | R<sup>2</sup> | Adjusted R<sup>2</sup> | p-val | S<sub>e</sub> |
| --- | --- | --- | --- | --- | --- |
| **GM Model- Step 1** |  |  |  |  |  |
| *High Beta FC* | 0.539 | 0.291 | 0.282 | <0.001 | 2.39 |
| **GM Model- Step 2** |  |  |  |  |  |
| *High Beta FC* | 0.538 | 0.361 | 0.345 | <0.001 | 2.28 |
| *Left temporal lobe ctx* | -0.265 |  |  |  |  |

*Note.* The values of the $\beta$ coefficient, p-value, $r^2$ , adjusted $r^2$ coefficients and the standard error of the estimation are presented for each of the models and their respective steps.

### 3.3 Behavioural implications of alcohol-related networks

To explore the association of the FC networks related to future alcohol consumption with the behavioural variables, a CBPT approach based on spearman correlations were carried out. We analyzed the correlation between the PLV values of the different networks and the scores obtained in the SSS-V, BIS-11 and the executive component. As a result, it was found that within the alpha network there was a group of frontoparietal links with significant and positive correlation (*p* = 0.03, rho = 0.29) with the executive component (Figure 7A). At the same time, within the alpha network we found a left-lateralized cluster of links between frontal, temporal and parietal regions that presented a significant and positive correlation (p-value = 0.02, rho = 0.26) with the SSS-V scores (Figure 7B). Finally, no significant correlations were found between any of the analyzed networks and the BIS-11 questionaries. After controlling for volume conduction and source leakage, all extracted links remained below the significance level.

**Figure 7.**
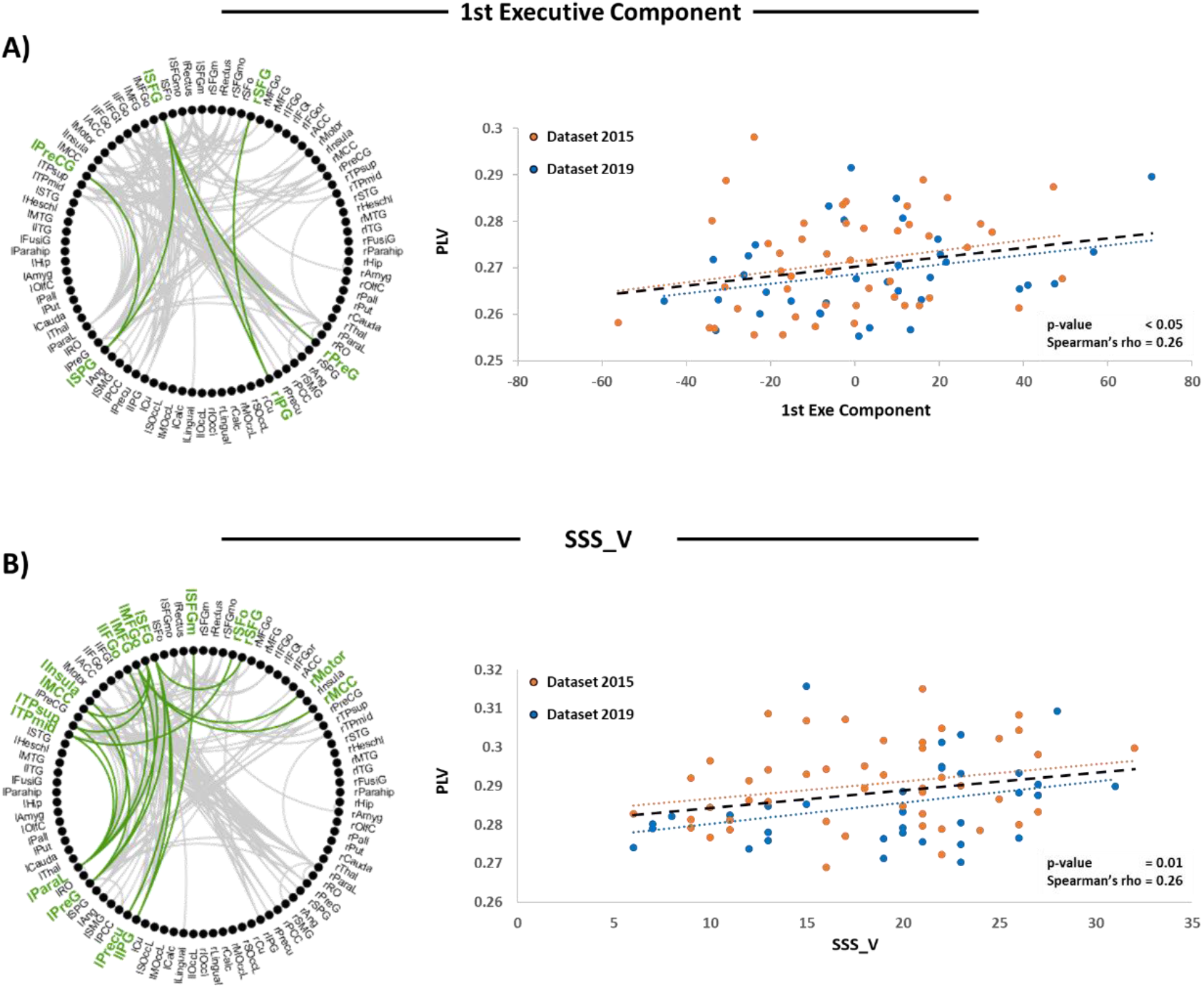
CBPT analysis between alpha network and the behavioural variables. A: On the left is represented in green the cortical organization of the cluster obtained after the analysis and in grey the analyzed alpha network. On the right is the correlation between the functional connectivity of the links marked in green and the value of the executive component. B: On the left is represented in green the cortical organization of the cluster obtained after the analysis and in grey the analyzed alpha network. On the right is the correlation between the functional connectivity of the links marked in green and the value of the SSS-V_V questionnaire.

## 4 Discussion

Current study investigated the connection between the physiological condition of the brain in 14-year-old adolescents and their future alcohol consumption patterns. As a result, we obtained that those adolescents who later engaged in more significant alcohol consumption displayed heightened connectivity values across all frequency bands. Once described, these functional networks were used as potential predictors of consumption together with neuroanatomical and behavioural variables to elaborate a comprehensive predictive model using multiple stepwise linear regressions. The findings highlighted that high beta band functional connectivity and the volume of the left temporal cortex functioned as main predictors of subsequent alcohol consumption. Finally, in a post-hoc analysis, the study illustrated the association between specific functional connections in the brain and the levels of sensation-seeking and executive function in adolescents.

### 4.1 Neurobiological and behavioural state of future alcohol drinkers

Prospective analysis of functional brain status with respect to adolescent consumption has been scarcely addressed in the literature. Most of the studies approached to this issue from functional magnetic resonance imaging (fMRI), finding that future alcohol and substance abuse is associated to lower local activation of frontal, parietal and subcortical regions during the performance of inhibition tasks ^35,36^ and resting state ^15^. However, holistic paradigms that consider the interaction of different brain regions, known as connectomics, have great potential for describing and studying human behaviour ^37^. In this sense, functional connectivity studies evaluated with fMRI have shown alterations between frontal regions and subcortical nuclei ^38^ and within frontoparietal connections ^19^ in adolescents prone to alcohol consumption. Nevertheless, the use of electrophysiological techniques facilitates the characterization of cortical brain dynamics with exceptional temporal resolution ^39^, which allows us to obtain comprehensive information on the underlying neurofunctional mechanisms associated with substance misuse. To our knowledge, only a couple of studies employed electrophysiological measures of FC in longitudinal studies in order to establish predictors of early alcohol consumption ^21,40^. Our study builds upon prior research by revealing links between hyperconnectivity patterns and future alcohol consumption that involve the entire frequency spectrum analyzed. Interestingly, this hyperconnectivity traits showed significant correlations regardless of the sample analyzed, indicating the consistency of these results. Furthermore, the existence of such differences before the onset of drinking habits should invite us to review previously obtained results on the relationship between intensive consumption and connectivity patterns ^8,41^. Specifically, in a study conducted by Correas et al., (2015) 18-year-old heavy alcohol consumers showed a reduction in functional connectivity in the alpha band, but an increase in the theta and beta bands ^8^. Our findings suggest that these variations in theta and beta could be traits predating alcohol use while decreased alpha connectivity in later heavy drinkers might indicate alcohol’s adverse effects on neurodevelopment. Nonetheless, more extensive longitudinal studies are required to unravel the intricate relationships between neural predispositions to heavy alcohol consumption, its subsequent effects, and its interplay with standard neurodevelopment during the transition from childhood to adulthood.

Regarding structural neuroimaging, recent longitudinal studies suggest that certain volumetric brain differences linked to alcohol consumption can be observed before individuals begin drinking ^13,14,42^. Consistent with these results, the present research found that a smaller left temporal cortex, caudate, and cerebellar white matter volume, along with larger right temporal volume and frontal white matter, were predictors of future alcohol habits. Nevertheless, contrary to previous reports ^14^, the right temporal cortex showed a positive relationship with consumption. In addition, differences observed by other authors within the frontal grey matter ^42^, the cingulum ^43^ or nucleus accumbens ^14^ were not statistically significant in our sample. Contrary to the discouraging results, the collective conclusion underlines the predictive value of the variables derived from structural magnetic resonance imaging and their complementarity to the functional variables extracted from electrophysiology.

Finally, the study of behavioural traits related to substances use predisposition have reported associations between poorer executive behaviour ^11^, impulsivity and sensation seeking ^5^ with future consumption. However, such associations were not significant across our sample, probably due to the fact that self-report questionnaires could be not robust enough to consistently reflect the cognitive status of adolescents ^44^. Nevertheless, post-hoc analysis revealed significant correlations with sensation-seeking and dis-executive behaviour. Consistent with prior findings, dis-executive behaviour correlated with functional connectivity in alpha band within the central executive network ^45^. Conversely, sensation-seeking was significantly linked to functional connectivity involving multiple interhemispheric frontal and left frontotemporal connections. Such observations concur with past studies, highlighting connectivity between frontal regions and substance-seeking behaviours in substance-abusing adults ^46^. Despite prior evidence linking impulsivity scales to functional connectivity ^47^, our analysis found no significant correlations. However, it’s notable that many impulsivity-related networks involve cortico-subcortical connections, which our measurement system may not fully capture.

### 4.2 Predictive models in adolescent alcohol consumption and its implications

The existence of early biomarkers of alcohol consumption in adolescent electrophysiology is inherently informative. Nevertheless, the usefulness of such markers lies in their ability to predict consumption and thus provide us the opportunity to establish prevention strategies. To date, the most in-depth study on predictors of adolescent alcohol consumption was conducted by Squeglia et al., 2017. This prospective study employed neuroimaging, sociodemographic, and behavioral variables in a random forest analysis. Aligned with the previously mentioned results, this study concluded that reduced cortical thickness in certain brain regions and altered activity in areas like the cingulate or precuneus during cognitive tasks were significant predictors of future alcohol use. However, this research did not include measures of electrophysiology that could contribute significantly to the description of consumption-prone profiles. Consequently, through this work we aimed to group the findings into behavioural, structural and functional connectivity modalities in regression models capable of describing the predictive potential of these variables. As a result, the regression analysis was able to build a model that explained 51% of the variance of future alcohol consumption with a root squared mean error of ± 2.61 SAUs where functional connectivity in the beta band and the volume of left temporal cortex were the main predictors, reflecting the usefulness of the above-mentioned measures.

Within the framework of neurobiological differences predisposing to consumption, there is broad consensus in understanding these variations as the product of divergent trajectories in neuromaturational mechanisms during adolescence. In this line, numerous studies have hypothesized that the findings linked to heavy alcohol consumption, specifically reduced cortical volumes and diminished task-related BOLD responses, mirror characteristics of a precocious neuromaturational state ^35,36,48^. Once puberty begins, a cascade of hormonal signals triggers several changes throughout the brain that are intimately linked to the development of substance use risk profiles ^49–51^. Such neuromaturative changes includes cortical thinning, increase of white matter ^50,52,53^ and refinement of the dopaminergic ^44^ and GABAergic ^54^ neurotransmission. However, according to the dual systems theory ^55^, the changes do not occur simultaneously in all brain systems. In this way, at the onset of puberty, an early maturation of the systems involved in motivation and reinforcement mechanisms would be followed by a slower and gradual maturation of the cortical systems in charge of cognitive control ^56^. Then, the mismatch between the two systems would lead to an increase in risk-taking and sensation-seeking behaviours typical of this age group ^44^. In this scenario, given that the onset and development of puberty is highly individualized, the neurodevelopmental changes produced may be sufficiently different to explain the existence of consumption-prone profiles.

From an electrophysiological standpoint, some MEG studies have noted that during typical adolescent neuromaturation, there is an increase in FC within the theta, alpha, and beta bands, while a decrease is observed in the gamma band ^57,58^. Specifically, while the increase in theta connectivity was cortically distributed, the changes in the alpha and beta band were predominantly found in posterior regions ^58^. Considering these findings, the observed association between higher levels of alcohol consumption and changes in brain structure and function, such as reduced volumes in the temporal cortex and putamen, increased white matter volume, and enhanced functional connectivity, lends support to the hypothesis that accelerated cortical maturation may be linked to a heightened likelihood of heavy drinking. However, the concept of “pseudomaturation” becomes relevant when considering discrepancies in the expected neuromaturative patterns. Specifically, the cognitive profiles and certain neuroanatomical and neurofunctional markers indicative of future alcohol consumption identified in this study do not fit to the expectations associated with advanced stages of neurodevelopment. While our findings indicated elevated FC in prefrontal regions, such results seem at odds with standard electrophysiological maturation. These discrepancies might reflect altered prefrontal functional mechanisms, as observed in risk-taking adolescents ^59^, those susceptible to AUD ^60^, and in various psychopathologies that course with behavioural dysregulation^61^. On the other hand, our gamma band findings revealed a cortical distribution similar to those reported in the standard maturational trajectory, but in opposite direction ^57^. This frequency band deserves special interest since it has traditionally been associated with the functioning of interneurons ^62^, which are extensively remodelled during neurodevelopment and whose activity is necessary for the emergence of coordinated activity in the brain. Therefore, deviations in the maturation of GABAergic interneurons could induce cortical hyperexcitability, revealing itself as anomalous hyperconnectivity ^63^. Additionally, it is important to highlight that although greater connectivity could be related to advanced maturation profiles, and throughout neurodevelopment, an improvement in executive functions can be expected ^53^, in the present study we found an opposite relationship between these variables. These facts would highlight that the current findings go beyond the existence of maturational advances and reflect the presence of distinct neurodevelopmental profiles that influence the relationship between adolescents and their approach to alcohol. Considering the presence of various maturation profiles, the asynchrony between reward sensitivity and cognitive control systems, as well as the intrinsic development of each system, could be different among individuals. This would lead to the amplification of certain typical adolescent behaviours such as substance consumption. Moreover, the study of addictions currently accepts that there is no such thing as addictive personalities. Instead, there are certain risk traits that together with the contingencies of a given environment give rise to the onset and evolution of consumption ^64^.

### 4.3 Strengths and limitations

This study’s strengths include its experimental design, which enabled tracking multiple adolescents to examine variables related to consumption years before behavior onset. All participants underwent behavioural scales, MRI, and MEG recordings adhering to high-quality standards. The study also consistently observed associations between variables across two independent samples. However, limitations include scarce literature on electrophysiological measures for early markers of adolescent consumption, complicating comparisons beyond fMRI results. Additionally, variations in methodologies in M/EEG studies caution against direct comparisons. The study’s lack of fMRI data limits the examination of subcortical nuclei activity, crucial in neurodevelopment and alcohol consumption patterns. Lastly, the functional networks analyzed were exploratory and require validation through replication in an independent sample.

## 5 Conclusion

In summary, the results obtained throughout this study reflect the potential of functional connectivity variables in the study of adolescent drinking habits and behaviour. In addition, the creation of structural and functional models has the capacity to establish predictive models of alcohol consumption at early ages. Present work also indicates a strong relationship between neuromaturational courses and substance use. Thus, inter-individual differences in the timing and form of certain neurodevelopmental milestones could condition the relationship of adolescents with alcohol. These findings could be useful in the elaboration of preventive strategies at ages prior to the onset of consumption in which not only the risk markers mentioned above but also the social context in which young people grow up are taken into account.

## 6 Declaration of interest

The authors declare that the research was conducted in the absence of any commercial or financial relationships that could be construed as a potential conflict of interest.

## 7 Data availability

The data and code that support the findings of this study are available from the corresponding author, upon request.

## Notes

### Competing Interest Statement

The authors have declared no competing interest.

